# Extracellular Adenine Nucleotide and Adenosine Metabolism in Calcific Aortic Valve Disease

**DOI:** 10.1101/402685

**Authors:** Barbara Kutryb-Zajac, Patrycja Jablonska, Marcin Serocki, Alicja Bulinska, Paulina Mierzejewska, Daniela Friebe, Christina Alter, Agnieszka Jasztal, Romuald Lango, Jan Rogowski, Rafal Bartoszewski, Ewa M. Slominska, Stefan Chlopicki, Jürgen Schrader, Magdi H. Yacoub, Ryszard T. Smolenski

## Abstract

Extracellular nucleotide catabolism contributes to immunomodulation, cell differentiation and tissue mineralization by controlling nucleotide and adenosine concentrations and its purinergic effects. Disturbances of purinergic signaling in valves may lead to its calcification. This study aimed to investigate the side-specific changes in extracellular nucleotide and adenosine metabolism in the aortic valve during calcific aortic valve disease (CAVD) and to identify the individual enzymes that are involved in these pathways as well as their cellular origin.

Stenotic aortic valves were characterized by reduced levels of extracellular ATP removal and impaired production of adenosine. Respectively, already reduced levels of extracellular adenosine were immediately degraded further due to the elevated rate of adenosine deamination. For the first time, we revealed that this metabolic pattern was observed only on the fibrosa surface of stenotic valve that is consistent with the mineral deposition on the aortic side of the valve. Furthermore, we demonstrated that non-stenotic valves expressed mostly ecto-nucleoside triphosphate diphosphohydrolase 1 (eNTPD1) and ecto-5’nucleotidase (e5NT), while stenotic valves ecto-nucleotide pyrophosphatase/ phosphodiesterase 1, alkaline phosphatase and ecto-adenosine deaminase (eADA). On the surface of endothelial cells, isolated from non-stenotic valves, high activities of eNTPD1 and e5NT were found. Whereas, in valvular interstitial cells, eNPP1 activity was also detected. Stenotic valve immune infiltrate was an additional source of eADA. We demonstrated the presence of A1, A2a and A2b adenosine receptors in both, non-stenotic and stenotic valves with diminished expression of A2a and A2b in the former.

Extracellular nucleotide and adenosine metabolism that involves complex ecto-enzyme pathways and adenosine receptor signaling were adversely modified in CAVD. In particular, diminished activities of eNTPD1 and e5NT with the increase in eADA that originated from valvular endothelial and interstitial cells as well as from immune inflitrate may affect aortic valve extracellular nucleotide concentrations to favor a proinflammatory milieu, highlighting a potential mechanism and target for CAVD therapy.

## 1. Introduction

Calcific aortic valve disease (CAVD) is a slow progressive disorder related to degeneration and mineralization of valve leaflets. (1) Increased stiffness of the leaflets results in limited valve opening and leads to heamodynamic overload on the left ventricle, followed by a valvular cardiomyopathy. (2) Currently, no medical therapies are available to prevent the development and progression of CAVD. Surgical aortic valve replacement (AVR) or transcatheter aortic valve implantation (TAVI) remain the only effective and durable methods for treatment of severe aortic stenosis. In the USA, CAVD is the primary cause of 95,000 valve replacements performed annually and the number of these operations have steadily been increasing over the last few decades. (3) In early stages, CAVD is an active cell-regulated process initiated by endothelial disruption with macrophages and T cell infiltration with accumulation and oxidation of lipoproteins. (4) These factors activate quiescent valvular interstitial cells (qVIC) to activated VICs (aVIC), which are characterized by the expression of smooth muscle cells alpha actin (α-SMA). Activation of VIC is associated with increased extracellular matrix production and remodeling as well as expression of matrix metalloproteinases and secretion of proinflammatory cytokines, which all together result in pathological fibrosis and chronic inflammation of the valve. (5) Simultaneously, in the presence of proteins associated with chondro- and osteogenesis, like osteopontin or after stimulation by oxidized LDL and cytokines (TGF-β), VICs undergo osteoblastic differentiation (obVIC), which is a direct cause of aortic valve mineralization. (6) Another potent regulators of osteoblastic VIC differentiation is extracellular adenosine triphosphate (ATP), adenine nucleotide that acts by purinergic P2 receptors, and its breakdown product, adenosine that triggers cell-signaling effects by the activation of P1 receptors. (7,8) It has been indicated that nucleotides and their catabolites play a relevant role in the extracellular compartment, besides an undeniable function within cell. In the cardiovascular system, ATP and ADP (adenosine diphosphate) are released by various cells, after their stimulation by shear stress, hypoxia, hyperglycemia, inflammation or platelets activators. (9) Despite much lower concentration of ATP in extracellular space (nanomolar) than in cell (milimolar), its role as a signaling molecule seems to be important since it is known that nucleotides exist in the pericellular space at micromolar levels. (10) Moreover, activation of nucleotide receptors not only induces osteoblastic differentiation, but also affects: 1) inflammation by stimulation of adhesiveness and transmigration of immune cells through endothelium layer, 2) oxidation of LDL by controlling superoxide production, 3) thromboregulation by platelets activation or 4) bone remodeling by osteoclasts activation and increase of RANKL expression. (11–14)

Under normal physiological conditions extracellular nucleotides are inactivated through hydrolysis by cell-surface ecto-enzymes. (15) The first enzyme engaged in this cascade is an ecto-nucleoside triphosphate diphosphohydrolase 1 (eNTPD, CD39), which converts ATP to ADP and then to AMP (adenosine monophosphate). (16) AMP is rapidly hydrolyzed by ecto-5’nucleotidase (e5NT, CD73) to form adenosine, which is degraded by the last enzyme of this pathway, ecto-adenosine deaminase (eADA) that is fixed to the membrane by CD26 protein and/or adenosine receptors. (17,18) Extracellular nucleotides may also be catabolized by other enzymes such as ecto-nucleotide pyrophosphatases/ phosphodiesterases (eNPPs) or alkaline phosphatase (ALP). (19,20) In turn, upstream pathways, which lead to ATP synthesis from AMP seems to be the least important, because of the minimal ecto-kinase activity. (21) Except for the removal of nucleotides from the extracellular space, the significant function of ecto-nucleotidases is the production of adenosine, which attenuates inflammation, lipid oxidation, platelets reactivity and bone resorption. (22–25) Thus, the pericellular concentration of nucleotides and adenosine are strictly dependent on the production and breakdown of these molecules. Despite a few reports of selective changes in ecto-nucleotidase activities in CAVD, these analyses were limited in scope and there is no overall assessment of extracellular pathways of nucleotide and adenosine metabolism in the aortic valve.

Therefore, this study aimed to comprehensively examine extracellular nucleotide and adenosine metabolism in the human aortic valve and calcific aortic valve disease. For the first time, we have investigated the total flux between nucleotide degradation, adenosine production and its breakdown on both surfaces of stenotic and nonstenotic aortic valves. Moreover, we have identified individual enzymes responsible for these pathways as well as we have indicated their cellular origin.

## 2. Methods

### 2.1 Patients and tissue collection

The study was performed based on the standards of the Declaration of Helsinki. The study was approved by the local ethical committee and informed consent has been obtained from the patients. Stenotic aortic valves were collected during valve replacement for CAVD (total n=61, mean age: 60, median age 62; 40 males; 18 females; age range: 36-74), while non-stenotic aortic valves (total n=34, mean age: 53, median age: 53; 22 males; 13 females; age range: 28-75) were obtained during heart transplantation or Bentall procedures. Clinical characteristics of aortic valve stenosis patients and control patients included for the analysis of nucleotide and adenosine degradation rates on the fibrosa surface of the valve **(Figure 1)**, as well as their comorbidities, pharmacotherapy and laboratory data, are described in **Table 1**. For other analyses, a smaller groups of patients were used as indicated in the figure legend. Dissected human aortic valve leaflets were immediately placed into ice-cold physiologic salt solution and transported to the laboratory on ice within 30 min of harvest.

**Table 1.** Patient characteristics. Clinical characteristics of control patients and aortic valve stenosis patients included for the analysis of nucleotide and adenosine degradation rates on the fibrosa surface of the valve (Figure 1). Results are shown as mean ± SEM or percentage; *p<0.05 vs. control group.

|  | Control<br>(n=24) | Aortic stenosis<br>(n=52) |
| --- | --- | --- |
| Age, yrs | 53 ± 3 | 60 ± 1 |
| Female/Male | 6/18 (25/75%) | 16/36 (31/69%) |
| Body weight [kg] | 80 ± 3.0 | 83 ± 2.5 |
| LIPID PROFILE |  |  |
| Total cholesterol [mg/dl] | 161.9 ± 7.9 | 183.9 ± 7.9 |
| Low density lipoproteins [mg/dl] | 96.3 ± 12.0 | 111.5 ± 7.5 |
| Triacylglycerols [mg/dl] | 116.9 ± 7.9 | 127.7 ± 7.9 |
| High density lipoproteins [mg/dl] | 42.3 ± 3.1 | 44.7 ± 1.9 |
| GLYCEMIA |  |  |
| Fasting glucose [mg/dl] | 104.6 ± 4.7 | 110.7 ± 1.1 |
| Glycated hemoglobin HbA1c [%] | 5.6 ± 0.08 | 6.1 ± 0.16* |
| COAGULATION PARAMETERS |  |  |
| Prothrombin time [s] | 12.2 ± 0.31 | 12.0 ± 0.13 |
| International Normalized Ratio | 1.05 ± 0.02 | 1.06 ± 0.01 |
| Fibrinogen [g/l] | 3.55 ± 0.03 | 4.02 ± 0.15* |
| BLOOD PRESSURE |  |  |
| Systolic pressure [mm Hg] | 129 ± 6.3 | 131 ± 2.7 |
| Diastolic pressure [mm Hg] | 73 ± 2.2 | 76 ± 1.9 |
| COMORBIDITIES |  |  |
| Aortic regurgitation | 0 (0%) | 52 (100%) |
| Aortic insufficiency | 20 (83%) | 18 (35%) |
| Aortic aneurysm | 13 (54%) | 15 (53%) |
| Hypertension | 11 (46%) | 37 (71%) |
| Coronary artery disease | 4 (15%) | 18 (35%) |
| Hyperlipidemia | 6 (33%) | 26 (50%) |
| Diabetes mellitus | 2 (8%) | 17 (33%) |
| PHARMACOTHERAPY |  |  |
| Antihypertensives | 10 (19%) | 36 (69%) |
| Statins | 4 (15%) | 26 (50%) |
| Antithrombotics | 11 (45%) | 27 (52%) |
| Antidiabetics | 2 (8%) | 15 (29%) |

**Figure 1.**
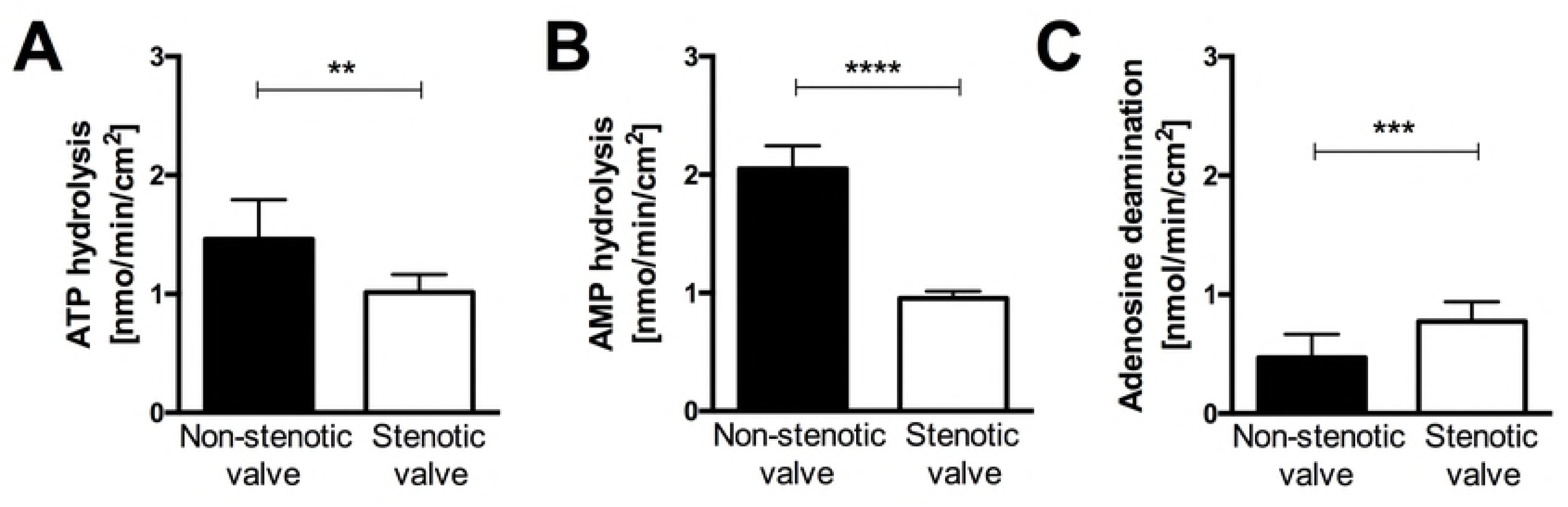
In aortic stenosis, activities of nucleotide-degrading ecto-enzymes are decreased, while adenosine catabolism is increased on the fibrosa surface of aortic valve. Rates of ATP hydrolysis **(A)**, AMP hydrolysis **(B)** and adenosine deamination **(C)** on the fibrosa surface of non-stenotic (*n*=24) and stenotic (*n*=52) aortic valve. The average rate of nucleotide or adenosine convertion for each valve was estimated from measurements for three leaflets independently, in the sites free of calcificaction. Results are shown as mean ± SEM; \*\**p*<0.01; \*\*\**p*<0.001; \*\*\*\**p*<0.0001 vs. non-stenotic valve by Mann-Whitney test.

### 2.2 Determination of aortic valve surface ecto-enzymes activities

For the determination of ecto-enzymes activities, valve leaflets were weighed and washed in Hanks Balanced Salt Solution (HBSS). Then, intact aortic valve leaflets were divided into 0.2 cm^2^ sections and directly placed into incubation solution. The modified assay system based on exposition into incubation medium by fibrosa and ventricularis surfaces separately. A valve leaflet fragment was fixed under the 0.5 cm diameter hole drilled in the bottom of one well of 24-well plate. It was supported by a plastic plate and the pressure was adjusted to ensure an effective seal. The leaflet fragment, clamped between two plastic plates, fully sealed the area exposed to the incubation medium. (26) Next, each well has been washed twice with HBSS and 1 ml of HBBS with 50 μM adenosine, ATP or AMP was sequentially added with medium exchange after each substrate. 5 μM erythro-9-(2-hydroxy-3-nonyl)adenine (EHNA), an inhibitor of adenosine deaminase, was present during incubation with ATP and AMP to block the conversion of adenosine to inosine. (27) Although, nucleotides and nucleosides are maintained in extracellular space at nanomolar level, we adjusted the substrate concentration to micromolar as these compounds operate on the cell surface in the “pericellular halo”. (18) To ensure that evaluated activities originate exclusively from the action of extracellular enzymes, part of experiments were conducted with the nucleoside transport inhibitor: 5 μM S-(4-Nitrobenzyl)-6-thioinosine (NBTI). (28)

After 0, 5, 15 and 30 min of incubation at 37°C samples were collected and concentrations of nucleotides and nucleosides were measured by reversed-phase HPLC according to the method described earlier. (29) Enzyme activities were calculated from linear phase of the reaction and in the main experiment, the rates were normalized to the surface area. Final results for each patient based on the average activity obtained from 3 valve leaflets. After the experiment, valve leaflet fragments were washed in HBSS, dried and frozen at −80° C for later use.

### 2.3 Determination of valve deposits compounds concentrations

Sections of aortic valve leaflets, previously used for the estimation of ecto-enzymes activities, were quickly thawed and dissolved separately in 6 M HCl at 95°C for 24 h followed by centrifugation at 2000 x *g* during 30 min. The supernatant was collected and diluted with deionized water and used for the determination of calcium and magnesium or diluted with 0.6 M H_2_SO_4_ for phosphate determination. Calcium content was estimated by Arsenazo III method, which relies on the formation of blue-purple complex at neutral pH. (30) Valvular magnesium concentration was analyzed using Calmagite, which forms a red complex with Mg^2+^ in an alkaline solution. (31) In turn, phosphate content was determined through a production of the green complex with malachite green molybdate under acidic conditions. (32) The intensity of solution decoloration was measured spectrophotometrically (Microplate Spectrophotometer Synergy HT, BioTek Instruments, Inc., Winoosk, VT) at 630 nm for Ca^2+^ and PO ^3-^ and 490 nm for Mg^2+^. Results were expressed as mg of calcium, magnesium or phosphate per wet weight of tissue (mg/g wt).

### 2.4 Histological analysis

Representative non-stenotic (n=4) and stenotic (n=3) aortic valve leaflets were fixed in 4% buffered formaldehyde, decalcified (if necessary) and embedded in paraffin. Then, the paraffin-embedded aortic valve leaflets were cut into 5 μm-thick cross-sections using a histological microtome, placed on microscopic slides and deparaffinized prior to staining. Sections were stained with hematoxylin and eosin (HE) for general morphology. For the assessment of specific aortic valve morphology, adjacent sections were stained according to Masson’s Trichrome (TR) standard protocol and Orcein Martius Scarlet Blue (OMSB) protocol. (33) These stainings allowed to characterize non-stenotic and stenotic valve composition, including cellular components as well as extracellular matrix fibers (loose connective tissue), collagen fibers (dense connective tissue), calcium nodules and myofibroblast-like cells, which far exceeds the capabilities of standard staining for calcium deposits. Images were acquired using a Dot Slide automatic scanning station (Olympus, Japan), stored as tiff files and analyzed automatically by the Image Browser software (Carl Zeiss). Areas of calcification were assessed in 6 cross-sections per each valve stained with TR and OMSB. Data were shown as the mean area of calcification expressed as the percentage of total aortic valve area.

### 2.5 Imunofluorescence analysis

Adjacent tissue sections to sections used for histological stainings were used to immunofluorescence analysis (IF). 5 μm-thick paraffin-embedded aortic valve cross-sections were collected on polylysine-covered microscopic slides and deparaffinized using a standard protocol. Next, sections were pretreated according to the citrate-base HIER (*Sigma*) protocol to unmask the antigens and epitopes in formalin-fixed and paraffin-embedded sections. Human primary aortic valve endothelial (hVEC) and interstitial (hVIC) cells intended to IF were seeded on 96-well optical-bottom plate (*Nunc ThermoFisher, USA*) at a density 1 × 10^4^ cells/ well in a total volume of 200 uL cell culture medium. 24 hours after seeding of hVEC and 72 hours after seeding of hVIC, cell culture medium was removed and rinsed 3 times with PBS. Cells were fixed using 100 uL 4% paraformaldehyde (pH 7.4) for 10 min at 37°C. Paraformaldehyde was removed and washed 3 times with PBS.

To reduce non-specific antibody binding, slices or cells were preincubated with a PAD solution (5% of normal goat serum and 2% of filtered dry milk) (*Sigma*). CD39 and CD73 were stained using mouse anti-human CD39 (*Novus*) and mouse anti-human CD73 (*Novus*) primary antibody, followed by a Cy3-conjugated goat anti-mouse secondary antibody (*JacksonImmuno*). eNPP1, ALP, ADA, vWF, CD26, CD45, A1R, A2aR, A2bR and A3R were stained using a rabbit anti-human eNPP1 (*Novus*), rabbit anti-human ALP (*Novus*), rabbit anti-human ADA (*Proteintech*), rabbit anti-human vWF (*Proteintech*), rabbit anti-human CD26 (*Genetex*), rabbit anti-human CD45 (*Genetex*), rabbit anti-human A1R (*Novus*), rabbit anti-human A2aR (*Novus*), rabbit anti-human A2bR (*Novus*), rabbit anti-human A3R primary antibody (*Novus*), followed by a Cy3-conjugated goat anti-rabbit secondary antibody (*JacksonImmuno*). Vimentin was stained using mouse anti-human vimentin conjugated with AlexaFluor488 (*Novus*). Primary antibodies were used at 1:100 final dilution (1h incubation), while secondary antibodies at 1:600 (30 min incubation). Negative controls omitted the primary antibodies (data not shown). Cell nuclei were counterstained with Hoechst 33258 (*Sigma*) (1:1500 final dilution, 5 min incubation). Images were recorded with an AxioCam MRc5 camera and an AxioObserved.D1 inverted fluorescent microscope (*Zeiss*) with appropriate filter cubes to show Cy3 (red), Alexa Fluor 488 (green) and Hoechst 33258 (blue) fluorescence, stored as tiff files and analyzed automatically using the Columbus Image Data Storage and Analysis System (Perkin Elmer). Total CD39, CD73, eNPP1, ALP, ADA, A1R, A2aR, A2bR and A3R positive area in aortic valves were measured in each slide and the percentage of total aortic valve cross-sectional area covered by red signal was calculated from six sections.

### 2.6 Gene expression

Human non-stenotic (n=6) and stenotic (n=9) aortic valves were directly lysed with QIAzol^®^ Lysis Reagent (*Qiagen, Hilden, Germany*) by shaking (5 min) in the presence of 3 mm diameter solid glass beads (*Sigma, USA*). Total RNA was isolated with RNeasy mini kit (*Qiagen*) according to the manufacturer’s instruction. To prevent DNA contamination, samples were pretreated with RNase-free DNase (*Qiagen*). The concentration of RNA was calculated based on the absorbance at 260 nm. RNA samples were stored at −70 °C until use. For the measurement of eNTPD1, e5NT, ADA, ADORA1, ADORA2a, ADORA2b and ADORA3 mRNA expression, TaqManOne-Step RT-PCR Master MixReagents (*Applied Biosystems, USA*) were used as described previously (34,35) according to the manufacturer’s protocol. The relative expressions were calculated using the comparative relative standard curve method. (36) We used housekeeping gene, TATA-binding protein (TBP), as the relative control. TaqMan probes ids used were: e5NT - Hs00159686_m1; eNTPD1 - Hs00969559_m1; ADA - Hs01110945_m1; ADORA1 Hs00181231_m1; ADORA2a - Hs00169123_m1; ADORA2b - : Hs00386497_m1; ADORA3 - Hs00252933_m1.

### 2.7 Non-stenotic aortic valve cells isolation and culture

Aortic valve endothelial (hVEC) and interstitial (hVIC) cells were isolated from non-stenotic human aortic valves (n=3) as was shown in **Figure S4**. The valve was digested with 5 mL collagenase A (0.15% w/v) for 10 min at 37°C to obtain hVEC. 5 mL of Endothelial Cell Growth Medium Medium (*Lonza, USA*) was added to stop the action of collagenase. To isolate hVIC, the valve was minced and further digested with 5 mL collagenase A (0.15% w/v) for additional 45 min at 37°C. 5 mL of DMEM (*Sigma, USA*) supplemented with 1 mmol/L L-glutamine, 10% FBS and 1% penicillin/streptomycin (v/v) (*Sigma, USA*) was added to neutralize the collagenase. Each of the suspensions, hVEC and hVIC, was purified using mesh filters 100 μm, 70 μm, 40 μm and centrifuged (150 × *g*, 4 min). After centrifugation, hVEC pellet was resuspended in EBM-2 Medium (*Lonza, USA*), while hVIC pellet in a standard Dulbecco’s Modified Eagle’s medium (DMEM, *Sigma, USA*) supplemented with 1 mmol/L L-glutamine, 10% FBS and 1% penicillin/streptomycin (v/v) (*Sigma, USA*). Cells were cultured at 37°C, in 5% CO_2_ atmosphere and used for experiments at passage 4.

### 2.8 Stenotic aortic valve cells isolation

Aortic valve endothelial and interstitial cells, as well as immune infiltrate, were also isolated from stenotic human aortic valves (n=3). hVEC and immune cells located in the upper layers of the valve were isolated after 10 min incubation with agitation in 5 mL of collagenase A (0.15% w/v) at 37°C. 5 mL of EBM-2 Medium (*Lonza, USA*) was added to stop the action of collagenase. Aortic valve transport medium and suspension obtained after the 1^st^ step of isolation were purified using mesh filters 100 μm, 70 μm, 40 μm. After centrifugation (150 × *g*, 4 min), pellets were resuspended in MACS buffer, pooled and used for FACS analysis. hVIC and immune cells derived from the deeper layers of the valve were isolated after mincing of the valve and additional digestion for 45 min in 5 mL of collagenase A (0.15% w/v) at 37°C. 5 mL of DMEM (*Sigma, USA*) supplemented with 1 mmol/L L-glutamine, 10% FBS and 1% penicillin/streptomycin (v/v) (*Sigma, USA*) was added to neutralize the collagenase. Petri place used for mincing of the tissue was washed using PBS and collected solution and suspension obtained after the 2^nd^ step of isolation were purified using mesh filters 100 μm, 70 μm, 40 μm. After centrifugation (150 × *g*, 4 min), pellets were resuspended in MACS buffer, pooled and used for FACS analysis.

### 2.9 Peripheral blood mononuclear cells isolation

Peripheral blood mononuclear cells (PBMC) were isolated from healthy adult donors (n=3) using a Histopaque procedure. Briefly, a layer of 3 mL Histopaque 1.077 g/mL (*Sigma, USA*) was applied to the layer of 3 mL Histopaque 1.119 g/mL (*Sigma, USA*) in a 15-mL conical centrifuge tube. Blood was applied to the Histopaque 1.077 g/mL layer and it was centrifuged at 400 × *g* for 30 min at room temperature. After centrifugation, the upper layer was aspirated to within 0.5 cm of the opaque interface containing mononuclear cells and discarded. The opaque interface was transferred into a clean conical centrifuge tube and washed with 0.9 % NaCl containing 5 mM EDTA. After centrifugation (250 × *g*, 10 min), the supernatant has been discarded and the cell pellet was resuspended with HBSS.

### 2.10 Determination of specific ecto-enzymes activities on the surface of aortic valve and immune cells

Human primary aortic valve endothelial and interstitial cells isolated from non-stenotic aortic valves were seeded on 24-well plates at a density of 0.05 × 10^6^ cells/well. Cells were used for the experiment at 90-100% confluency and washed with HBSS. Isolated human peripheral blood mononuclear cells and monocyte/macrophage cells (SC line, ATCC, cat. CRL-9855) were plated in 24-well cell culture plate at a density 0.2 × 10^6^ per well in a total volume of 1 mL HBSS. Cells were pre-incubated in HBSS for 15 min at 37°C with specific ecto-enzyme inhibitors, including 5 μM erythro-9-(2-hydroxy-3-nonyl) adenine for ADA1, 150 μM adenosine 5′-(α,β-methylene)diphosphate (AOPCP) for e5’ NT/CD73 (37), 500 μM levamisole hydrochloride for alkaline phosphatase (38), 150 μM 6-N,N-Diethyl-β-γ-dibromomethylene-D-adenosine-5′-triphosphate trisodium salt hydrate (ARL67156) for ecto-ATPases, mainly NTPDases (including eNTPD1/CD39) (39,40), 50 μM pyridoxal phosphate-6-azo(benzene-2,4-disulfonic acid) tetrasodium salt hydrate (PPADS) for ENPPs (41). After pre-incubation ecto-enzyme substrates were added, 50 μM adenosine AMP or ATP and cells were incubated at 37°C for 30 min. Samples of the incubation medium were collected in 0, 5, 15 and 30 min time points and analyzed for the concentration of nucleotides and their catabolites using HPLC as described above. Enzyme activities were calculated from linear phase of the reaction. The concentration of cell protein was determined using Bradford method according to the manufacturer’s instructions (*Bio-Rad, USA*).

### 2.11 Flow cytometry analysis

Cells were resuspended in MACS buffer, preincubated with FcR Blocking Reagent (*Miltenyi Biotech*) and stained with the following antibodies: anti-CD31-PE-Cy7, WM59 (*eBioscience*), anti-vimentin-AF488, RV203 (*Novus*), anti-bone sialoprotein-PE-Cy7 polyclonal (*Biorbyt*), α-SMA-eFluor660, 1A4 (eBioscience), anti-CD45-APC, 30-F11 (*BD Bioscience*), anti-CD4-PerCP-Cy5.5, RM405 (*eBioscience*), anti-CD8a-APC-H7, 53-6.7 (*BD Bioscience*), anti-CD19-PE-TR, SJ25-C1 (*LifeSpan BioSciences*), anti-CD11b-PE M1/70 (*BD Bioscience*), anti-CD14-AF488, M5E2 (*StemCell*), anti-CD73-FITC, 496406 (*R&D Systems*), anti-CD39-PE-Cy7, 24DMS1 (*eBioscience*), anti-CD26-PE 2A6 (*eBioscience*).

To identify aortic valve endothelial cells and individual subsets of aortic valve interstitial cells and immune cells we used a panel of antibodies against different cell-specific markers, including markers for endothelial cells (CD45-, CD31^high^), activated VIC (CD45-, Vim+, Sial-, αSMA^high^), activated/osteoblast-like VIC (CD45-, Vim+, Sial+, αSMA^int^), osteoblast-like VIC (CD45-, Vim+, Sial+, αSMA-), T helper cells (CD45+, CD8+), T cytotoxic cells (CD45+, CD4+), B cells (CD45+, CD19+), monocytes/macrophages (CD45+, CD11b+, CD14+) and granulocytes (CD45+, CD11b^int^, CD14-).

After 5 min of the incubation at room temperature cells were washed and resuspended in 200 μL MACS buffer for flow cytometry. Cell measurements were performed with a FACSCAnto II flow cytometer (BD Bioscience). For analysis, the placement of gates was based on fluorescence minus one (FMO) controls. The minimum number of events used to define a cell population was 150. The analysis was performed on individual aortic valves. The different cells subsets were enumerated and the percentage of CD39, CD73 and CD26 (adenosine deaminase binding-protein) and corresponding expression levels as measured by mean fluorescence intensity (MFI) was assessed.

### 2.12 Statistical analysis

Statistical analysis was performed using InStat software (GraphPad, San Diego, CA). Comparisons of mean values between groups were evaluated by one-way analysis of variance (ANOVA) followed by Holm–Sidak, or Sidak post hoc tests, two-way ANOVA followed by Sidak post hoc test, unpaired Student’s t-test, or Mann–Whitney U test, as appropriate. Normality was assessed using the Kolmogorov–Smirnov test, Shapiro–Wilk test, and the D’Agostino and Pearson Omnibus normality tests. The exact value of n was provided for each type of experiments. Statistical significance was assumed at *p*<0.05. Error bars indicated the standard error of the mean (SEM) unless otherwise described in the figure legend.

## 3. Results

### 3.1 Nucleotide and adenosine degradation rates on the surface of intact aortic valves

The activities of adenine nucleotide catabolism ecto-enzymes in intact non-stenotic **(Figure S1A-C)** and stenotic **(Figure S1D-F)** aortic valves were analyzed with two different assay methods. The use of a first method **(Figure S1A, S1D)**, which assumed the incubation of entire leaflet fragment in the substrate solution, resulted in a higher ATP hydrolysis, AMP hydrolysis and adenosine deamination than exposition only one side (aortic side, fibrosa) of the aortic valve leaflet (method 2, **Figure S1B, S1E**). Since the second method allows for the estimation of these activities on the fibrosa and ventricularis surfaces separately this method has been used in the later part of experiments. Nucleotide and adenosine degradation rates of intact non-stenotic aortic valves did not differ between fibrosa and ventricularis **(Figure S1G)**. In stenotic aortic valves, the rate of ATP and AMP hydrolysis was lower, while adenosine deamination was higher on the fibrosa than on ventricularis surface **(Figure S1H)**.

Comparing nucleotide and adenosine degradation rates on the fibrosa surface in a larger patients group that are characterized in **Table 1**, we observed lower ATP and AMP hydrolysis, as well as higher adenosine deamination, in stenotic aortic valve than in non-stenotic **(Figure 1)**. The blockade of a transmembrane nucleoside transport by NBTI did not affect the rate of product formation **(Figure S2)**.

Pre-operative echocardiographic parameters in a study group of patients **(Figure 2A)** exhibited severe aortic valve stenosis (aortic valve area < 1 cm^2^, aortic jet velocity > 4 m/s, mean gradient > 40 mmHg). Their valves, collected after aortic valve replacement, had a higher concentration of Ca^2+^, Mg^2+^, and PO_4_ ^3-^ than non-stenotic valves **(Figure 2B)**. Histological analysis of representative non-stenotic **(Figure 2C)** and stenotic **(Figure 2D)** aortic valves also revealed substantial areas of calcification in stenotic valves **(Figure 2E)**. However, the results obtained for nucleotide and adenosine degradation rates were determined on the stenotic valve surface in the areas free of calcification, as indicated for representative valves **(Figures S3A, S3B)**. We also exhibited that the rate of extracellular nucleotide metabolism did not differ between leaflets within the same non-stenotic **(Figure S3A)** and stenotic **(Figure S3B)** aortic valve.

**Figure 2.**
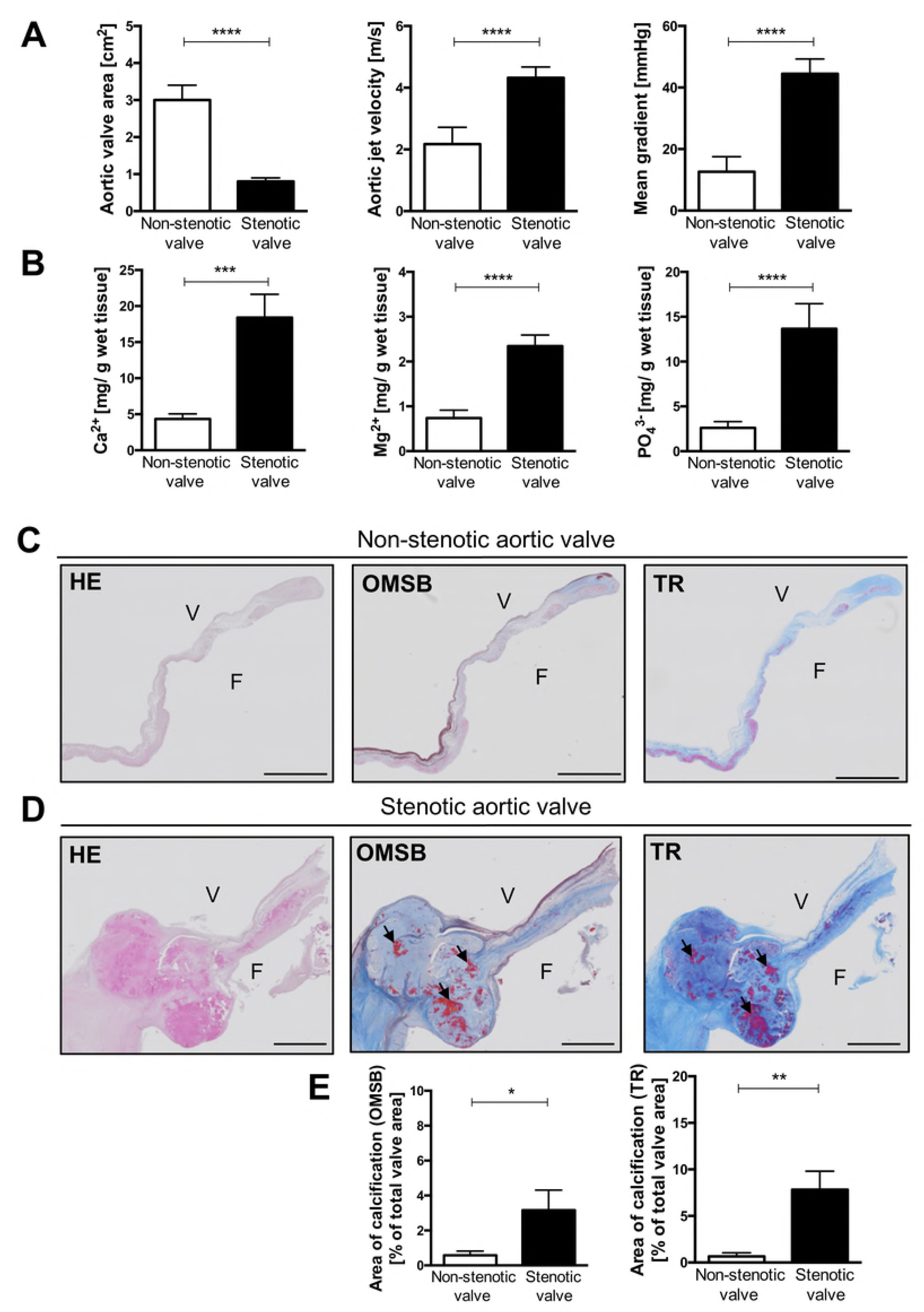
Characteristics of non-stenotic and stenotic aortic valves. Echocardiographic parameters **(A)** of patients before Bentall procedure (non-stenotic valves, *n*=24) and aortic valve replacement (stenotic valves, *n*=52). Concentration of Ca^2+^, Mg^2+^, PO ^3-^ **(B)** in non-stenotic (*n*=24) and stenotic (*n*=52) aortic valves. Representative images of non-stenotic **(C)** and stenotic **(D)** aortic valves stained with Hematoxilin and Eosin (HE). Orcein Mertius Scarlet Blue (OMSB) and Masson’s Trichrome (TR). F = fibrosa, V = ventricularis. Scale bar = 2 mm. HE staining was used for general microscopy. In OMSB staining cell nuclei were stain red, while purple/grey sections represent elastic fibers and elastic laminae, blue sections represent collagen fibers and red nodules represent calcium nodules. In TR staining, cell nuclei were stain dark pink/red, dark blue sections represent collagen fibers (dense connective tissue), light blue sections represent extracellular matrix fibers (loose connective tissue), purple nodules represent calcium nodules and red fibers represent myofibroblast-like cells. Calcium nodules were pointed by black arrows. Quantitative analysis of aortic valve calcification area **(E)** in non-stenotic (*n*=4) and stenotic (*n*=3) aortic valves. Results are shown as mean ± SEM; \**p*<0.05; \*\**p*<0.01; \*\*\**p*<0.001; \*\*\*\**p*<0.0001 vs. non-stenotic valve by Mann-Whitney test.

### 3.2 The presence of specific extracellular nucleotide and adenosine metabolism enzymes in aortic valves

At the next stage, we analyzed which enzymes involved in extracellular nucleotide and adenosine metabolism are found in aortic valves. For this purpose, microphotographs of histological stainings for representative aortic valves **(Figure 3A)** had been complied with immunofluorescence analysis **(Figure 3C)**. These results indicated that in both stenotic and non-stenotic aortic valves are enzymes that can be engaged in nucleotide and adenosine metabolism **(Figure 3B)**, including ecto-nucleoside triphosphate diphosphohydrolase 1 (eNTPD1, CD39), ecto-nucleotide pyrophosphatase/phosphodiesterase 1 (eNPP1), ecto-5’-nucleotidase (e5NT, CD73), alkaline phosphatase (ALP) and adenosine deaminase (ADA). Using the same fluorescence microscope settings, we determined the area of a specific signal for each enzyme **(Figure 3C)**. In a non-stenotic aortic valve, the most abundant signal area was observed for e5NT, eNTPD1, and eNPP1, while in stenotic valve for ALP and eNPP1 with a diminished signal area for e5NT and eNTPD1. Signal area for ADA was minor in both types of the valve, but it was directed towards a larger area in the stenotic valve. In turn, within the areas of calcification, we observed an accumulation of the signal for e5NT and ALP **(Figures S3C, S3D)**. However, total expression of e5NT, as well as eNTPD1, were lower in stenotic aortic valve than in non-stenotic **(Figure S3E)**.

**Figure 3.**
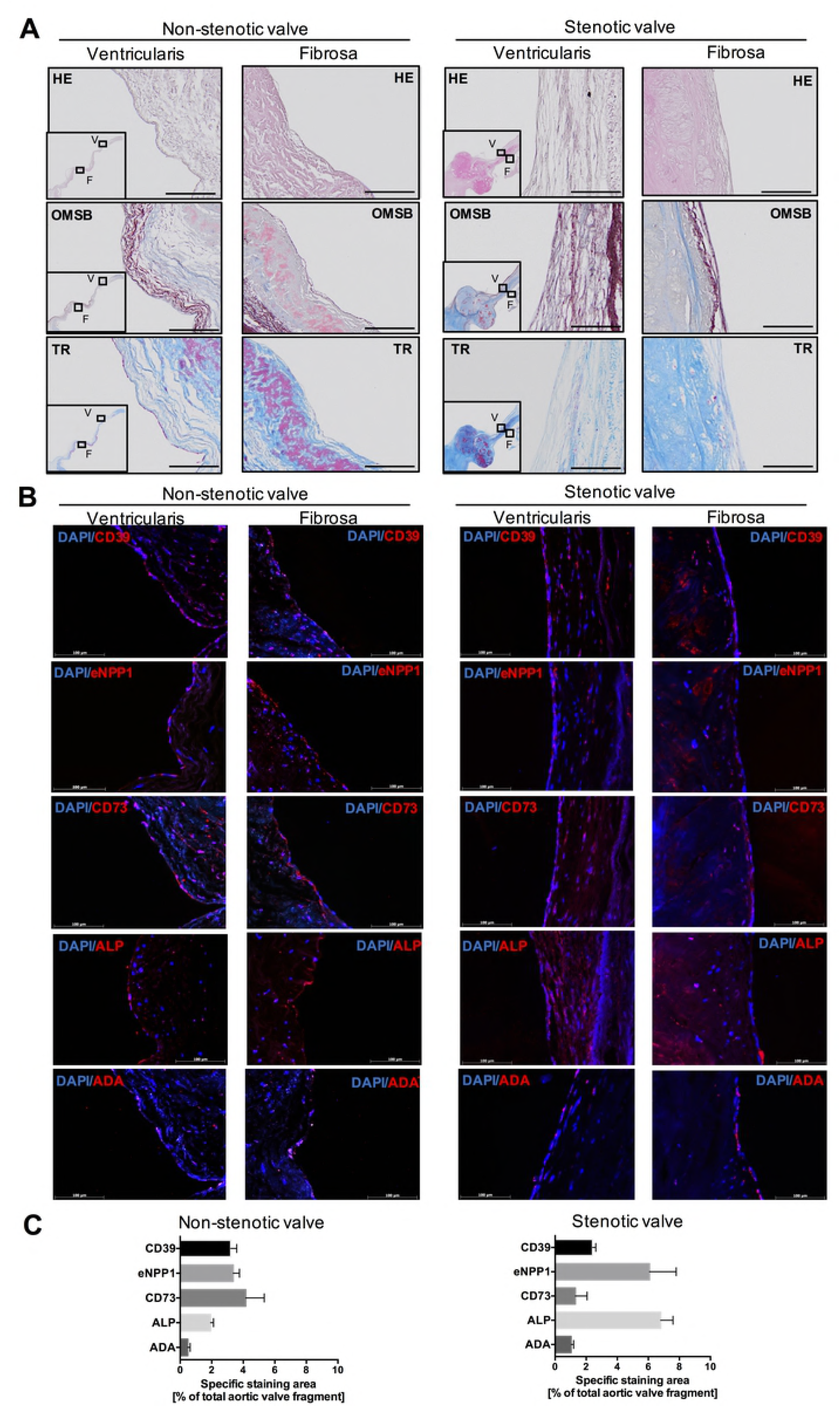
Human aortic valves, both non-stenotic and stenotic express nucleotide metabolism ecto-enzymes including ecto-nucleoside triphosphate diphosphohydrolase 1, ecto-nucleotide pyrophosphatase/ phosphodiesterase 1, ecto5’nucleotidase, alkaline phosphatase and adenosine deaminase. Representative images of fibrosa and ventriculatis of non-stenotic (*n*=4) and stenotic (*n*=3) aortic valves **(A)** stained with Hematoxilin and Eosin (HE), Orcein Mertius Scarlet Blue (OMSB) and Masson’s Trichrome (TR). Scale bar = 100 μm. Representative images of matching sections stained by immunofluorescence (red signal) for CD39 (ecto-nucleoside triphosphate diphosphohydrolase 1), eNPP1 (ecto-nucleotide pyrophosphatase/ phosphodiesterase 1), CD73 (ecto5’-nucleotidase), ALP (alkaline phosphatase) and ADA (adenosine deaminase). Quantitative analysis of CD39, eNPP1, CD73, ALP and ADA positive area **(C)** that corresponds to the specific signal for each enzyme. Fluorescence values of the negative control slices were substracted from the fluorescence value of the stained slices. Results are shown as mean ± SEM.

### 3.3 The activity of specific extracellular nucleotide and adenosine metabolism enzymes on non-stenotic aortic valve cells

Aortic valve endothelial and interstitial cells isolated from human non-stenotic aortic valves **(Figures 4A, S4A)** actively degraded nucleotides and adenosine on their surface **(Figure 4B-G)**. Using specific ecto-enzyme inhibitors, we observed that after incubation with ARL67156 (eNTPD1 inhibitor), about 70% of ATP hydrolysis was inhibited on both hVEC **(Figure 4B)** and hVIC **(Figure 4C)**. After the incubation with PPADS (eNPP1 inhibitor), we observed only about 10 % inhibition of ATP hydrolysis on hVEC **(Figure 4B)** and about 60% of inhibition on hVIC **(Figure 4C)**. Levamisole (ALP inhibitor) did not affect the ATP hydrolysis on hVEC **(Figure 4B)** but decreased its rate on hVIC about 20 % **(Figure 4C)**. The rate of AMP hydrolysis was decreased after addition of AOPCP (e5NT inhibitor) about 80 % on both, hVEC **(Figure 4D)** and hVIC **(Figure 4E)**, while levamisole did not affect AMP hydrolysis on both types of cells **(Figures 4D, 4E)**. EHNA (ADA1 inhibitor) almost completely abolished extracellular adenosine deamination on hVEC **(Figure 4F)** and hVIC **(Figure 4G)**.

**Figure 4.**
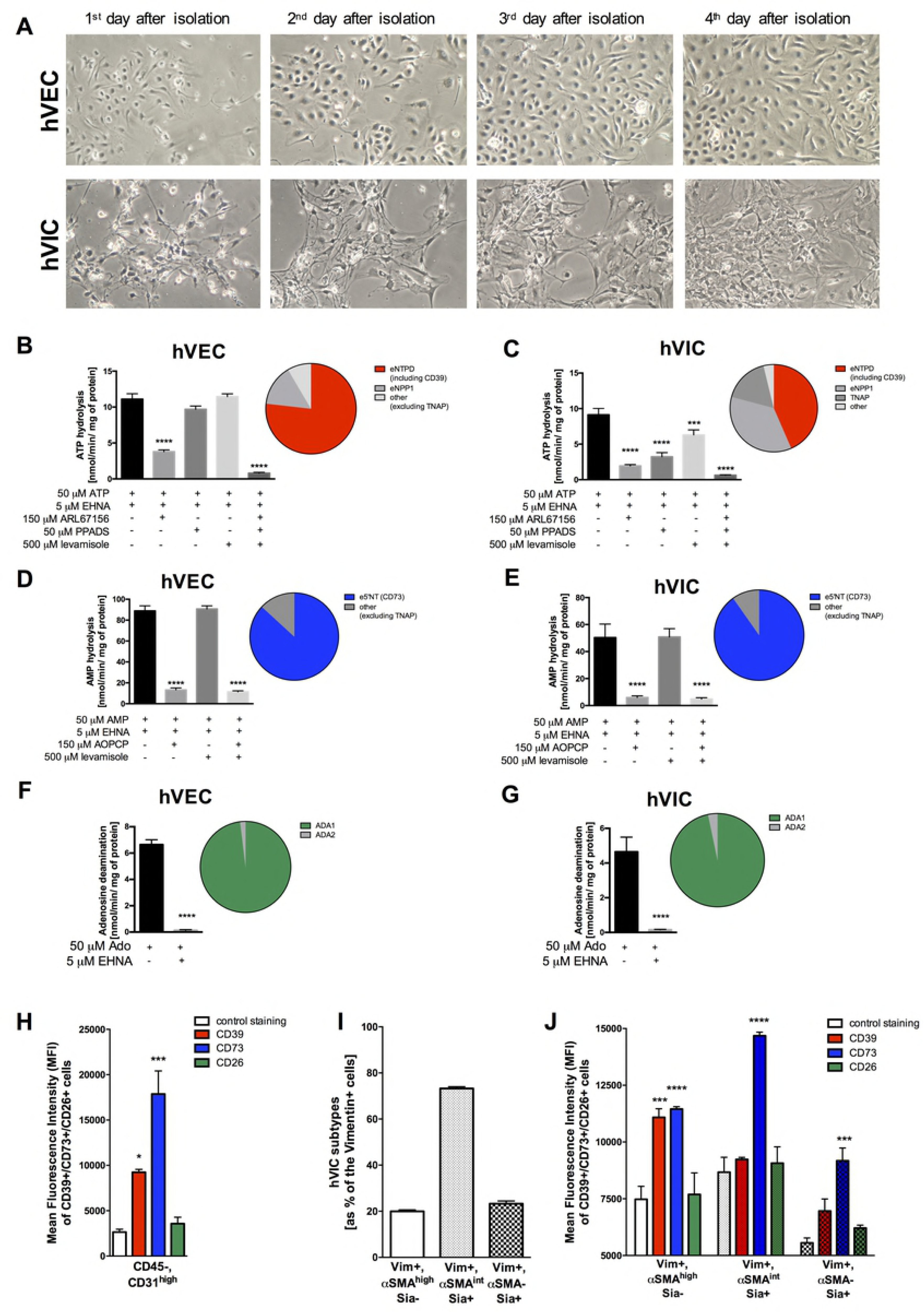
Aortic valve endothelial and interstitial cells are the main source of nucleotide-degrading ecto-nucleotidases. Representative images of cultured primary endothelial and interstitial cells isolated from human non-stenotic aortic valves in the following days after isolation **(A).** Magnification 100x. The rates of ATP hydrolysis **(B, C),** AMP hydrolysis **(D, E)** and adenosine deamination **(E, F)** on the surface of human aortic valve endothelial cells (hVEC; **B, D, F**) and interstitial cells (hVIC; **C, E, G**) in the presence of specific ecto-enzyme inhibitors. Flow cytometry analysis **(H-J)**. Mean fluorescence intensity of cell-surface CD39, CD73 or CD26 (ADA-binding protein) for CD31^high^ positive endothelial cells **(H)**. Percentage of interstitial cells (Vim+) as myofibroblast like-interstitial cells (αSMA^high^/Sia-), myo-/osteoblast-like interstitial cells (αSMA^int^/Sia+) and osteoblast-like interstitial cells (αSMA-/Sia+) **(I)** and mean fluorescence intensity of cell-surface CD39, CD73 or CD26 (ADA-binding protein) for each type of interstitial cells **(J)**. Results are shown as mean ± SEM; *n*=9 (independent isolations from 3 patients), \**p*<0.05, \*\*\**p*<0.001, \*\*\*\**p*<0.0001 vs. without specific inhibitors (**B-G)** or control staining **(H, J)** by one-way Anova followed by Holm-Sidak *post hoc* test **(B-E, H-J)** or student *t*-test (**F, G**).

### 3.3 The origin of individual enzymes of extracellular nucleotide and adenosine metabolism in stenotic aortic valve cells

As seen from the above results **(Figures 4A-G)**, eNTPD1, e5NT, and eADA1 (ecto-ADA1) are the main enzymes engaged in nucleotide and adenosine catabolism on non-stenotic aortic valve cells. Since the cultivation of the cells originated from stenotic valves is problematic, we isolated stenotic aortic valve endothelial and interstitial cells **(Figure S4B)** and immediately after isolation, we analyzed them with flow cytometry. As first, we compared the levels of these cell-surface proteins on stenotic aortic valve endothelial cells. The highest mean fluorescence intensity for CD31 positive cells expressed e5NT, then eNTPD1 and CD26 (ADA1-binding protein) **(Figure 4H)**. Also, the most of CD31 positive cells expressed on their surface e5NT, about 40 % expressed eNTPD1 and only about 10 % expressed CD26 **(Figure S4C)**. These results are in line with baseline levels of e5NT activity and signal for this protein that colocalized with vWF in immunofluorescence **(Figures 3B, S3D, S4C, S4C)**

After the isolation of stenotic aortic valve interstitial cells **(Figure S4B)**, which are vimentin positive (Vim+) cells, we gated them αSMA highly positive and bone sialoprotein (Sia) negative (Vim+, αSMA^high^, Sia-), Sia positive and αSMA intermediate positive (Vim+, αSMA ^int^, Sia+), Sia positive and αSMA negative (Vim+, αSMA-, Sia+) cells **(Figure 4I)**. The mean fluorescence intensity for e5NT was at the lowest level on the surface of Vim+, αSMA-, Sia+ cells **(Figure 4J)**. In turn, αSMA+ cells kept higher levels of e5NT on their surface **(Figure 4J)**. eNTPD1 was only detected on Siacells, which were highly positive for αSMA **(Figure 4J)**. Stenotic aortic valve interstitial cells exhibited almost undetectable levels of CD26 **(Figure 4J)**.

### 3.4 The origin and activity of individual enzymes of extracellular nucleotide and adenosine metabolism in immune cells

In a stenotic aortic valve, besides the valvular cells, there is also an inflammatory infiltrate **(Figures S5A, S5B)** that could be a source of nucleotide- and adenosine-degrading ecto-enzymes. Immune cells isolated from upper layers of the valve during the 1^st^ isolation step accounted for no more than 2 % of all cells, while immune cells isolated from deeper layers during the 2^nd^ isolation step were around 10 % of a total number of cells **(Figures S5C, 5A)**. In both cases, the dominant type of inflammatory cells were T helper cells (CD45+, CD4+), then B cells (CD45+, CD19+) and macrophages (CD45+, CD11b+, CD14-) **(Figure 5B)**. In contrast, isolates of inflammatory cells from all layers of the valve exhibited a small number of T cytotoxic cells (CD45+, CD8+) and granulocytes (CD45+, CD11b^int^, CD14-) **(Figure 5B)**. Immune cells were a poor source of e5NT, except a certain population of B cells **(Figures 5C, S5D)**. eNTPD1 originated mainly from B cells, monocytes/macrophages, and T helper cells **(Figures 5C, S5D)**. All populations of immune cells were an important source of CD26 **(Figures 5C, S5D)**.

**Figure 5.**
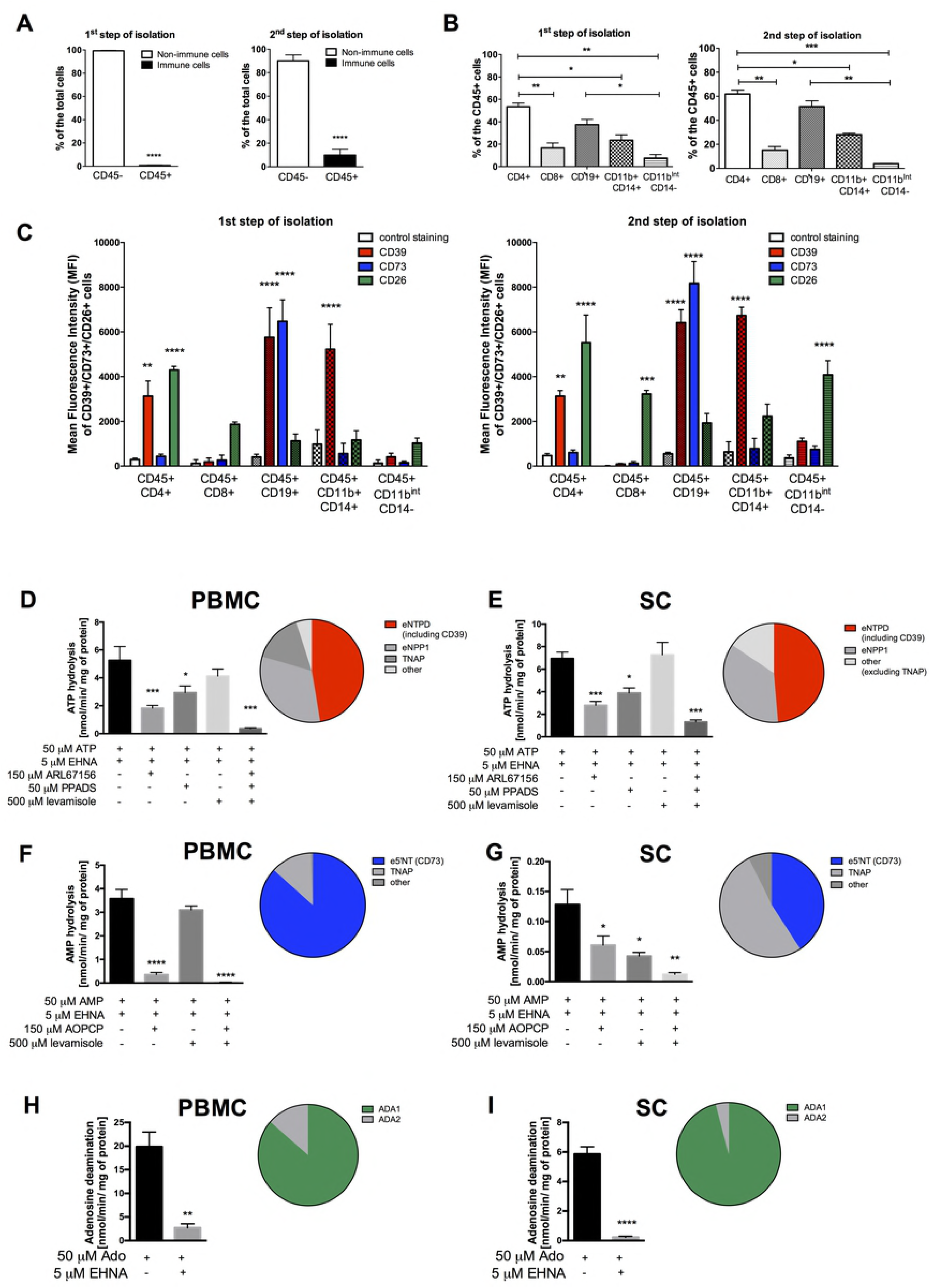
Stenotic aortic valve immune infiltrate is a smaller source of nucleotide-degrading ecto-nucleotidases but a larger of adenosine deaminase. Flow cytometry analysis of CD45 positive cells (immune cells) as a percentage of total isolated cells after 1^st^ step of isolation (cells located in the upper layers of the valve) and 2^nd^ step of isolation (cells located in the deeper layers of the valve) **(A)**. The composition of stenotic aortic valve immune infiltrate **(B)** expressed as a percentage (%) of total CD45+ cells, including T helper cells (CD45+,CD4+), T cytotoxic cells (CD45+,CD8+), B cells (CD45+,CD19+), monocytes/macrophages (CD45+,CD11b+, CD14+) and granulocytes (CD45+,CD11b^int^, CD14-). Mean fluorescence intensity of cell-surface CD39, CD73 or CD26 (ADA-binding protein) for each type of isolated immune cells **(C)**. Results are shown as mean ± SEM; *n*=9 (independent isolations from 3 patients), \**p*<0.05, \*\**p*<0.01, \*\*\**p*<0.001, \*\*\*\**p*<0.0001 vs. CD45- **(A)**, as indicated **(B)** or vs. control staining **(C)** by Student *t*-test **(A)**, one-way Anova followed by Tukey *post hoc* test **(B),** Holm-Sidak *post hoc* test **(C)**. The rates of ATP hydrolysis **(D, E),** AMP hydrolysis **(F, G)** and adenosine deamination **(H, I)** on the surface of human peripheral blood mononuclear cells (PBMC; **D, F, H**) and human monocyte/macrophages SC; **E, G, I**) in the presence of specific ecto-enzyme inhibitors. Results are shown as mean ± SEM; *n*=5-9, \**p*<0.05, \*\*\**p*<0.001, \*\*\*\**p*<0.0001 vs. without specific ecto-enzyme inhibitors **(D-I)** by one-way Anova followed by Holm-Sidak *post hoc* test **(D-G)** or Student *t*-test (**H, I**).

Despite that immune cells are not a dominant cell type in stenotic aortic valves, as we described above, they are still responsible for the origin of a certain pool of ecto-enzymes engaged in nucleotide and adenosine catabolism. As we have shown, the most significant in the number of cells and the presence of ecto-enzymes on their surfaces was the infiltrate of lymphocytes and monocytes/macrophages. Therefore during functional assays, we estimated the rates of nucleotide and adenosine degradation on the surface of human peripheral blood mononuclear cells (PBMC), which are mostly lymphocytes (42) and on monocytes/macrophages (SC cell line). We also used specific ecto-enzyme inhibitors to identify activities of individual enzymes. On the surface of lymphocytes, we observed 60 % of ATP hydrolysis inhibition after incubation with ARL67156, 30 % of ATP hydrolysis inhibition after incubation with PPADS, and only 10 % of ATP hydrolysis inhibition after using a levamisole **(Figure 5D)**. The effects of individual inhibitors on ATP hydrolysis was similar in monocytes/macrophages **(Figure 5E)**. In turn, AMP hydrolysis on lymphocytes was inhibited by 90 % after incubation with AOPCP and about 10 % after levamisole **(Figure 5F)**. The rate of AMP hydrolysis on monocytes/macrophages was inhibited by about 50-60 % after incubation with AOPCP as well as with levamisole **(Figure 5G)**. Adenosine deamination was inhibited by 80 % on lymphocytes and by 90% on monocytes/ macrophages after incubation with EHNA **(Figures 5H, 5I)**.

Comparing nucleotide and adenosine degradation rates on lymphocytes (PBMC) and monocytes/macrophages (SC cell line), the highest activity among all enzymes was observed for adenosine deaminase 1 (susceptible to inhibition by EHNA) on lymphocytes **(Figure 5H)**, while it was about 3.5 times lower on the surface of monocytes/macrophages **(Figure 5I)**. This is in line with our above results with high expression of an ADA-binding protein (CD26) on lymphocytes. The rate of ATP hydrolysis was at comparable levels on both types of cells **(Figures 5D, 5E)** and on monocytes/macrophages, it was similar to the rate of adenosine degradation **(Figures 5E, 5I)**. In turn, AMP hydrolysis was at the lowest level among all activities in both type of cells, while on the surface of monocytes/macrophages it was almost undetectable **(Figures 5F, 5G)**.

### 3.5 Adenosine receptors in human aortic valves

Since, ecto-enzymes engaged in nucleotide and adenosine metabolism play a key role in the bioavailability of adenosine in extracellular space for adenosine receptors, we determined which receptors are present in non-stenotic and stenotic aortic valves and which cells may be responsible for their origin. IF study **(Figures 6A, S6)** revealed that the most abundant among adenosine receptors in both non-stenotic and stenotic aortic valves was receptor A2a (A2aR) **(Figure 6B)**. A2b and A1 receptors (A2bR, A1R) occurred in smaller amounts **(Figure 6B)**. While A3 receptor (A3R) was not observed **(Figure 6B)**. Importantly, all three adenosine receptors that were found in aortic valves colocalized with endothelial cells **(Figure 6A)**. Whereas, their presence within deeper layers of the valve depended on the type of valve. A2aR was observed throughout the cross-section of non-stenotic valve **(Figure 6A)**, while it was almost undetectable in the deeper layers of stenotic valves and in calcifications **(Figures S6A, S6B)**. In contrast, A2bR was also observed in the depths of the stenotic valve, including calcification areas **(Figures S6A, S6B)**. Since, IF approach is not well adapted to conclude the differences in protein levels, we measured mRNA expression for adenosine receptors. This analysis confirmed their presence in aortic valves and revealed that expression of both, A2aR and A2bR was diminished in not calcified fragments of stenotic valves compared to non-stenotic **(Figures S6C)**.

**Figure 6.**
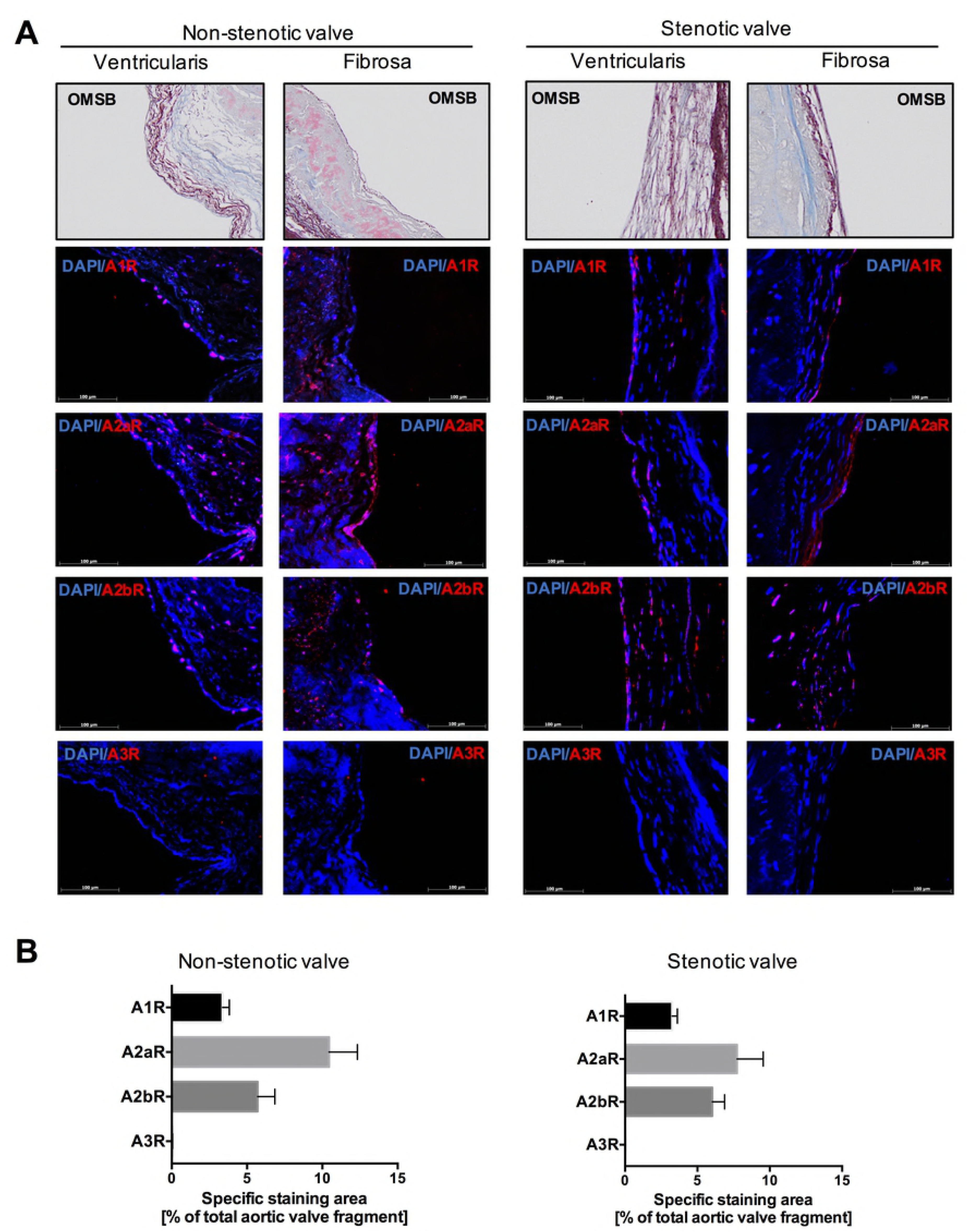
Adenosine receptors are widely express in human non-stenotic and stenotic aortic valves. Representative images of fibrosa and ventriculatis of non-stenotic and stenotic aortic valve (*n*=3) stained with Orcein Mertius Scarlet Blue (OMSB) and representative images of matching sections stained by immunofluorescence (red signal) for four types of adenosine receptors **(A)**. Scale bar = 100 μm. Quantitative analysis of A1R, A2aR, A2bR, A3R positive area that corresponds to the red signal **(B).** Fluorescence values of the negative control slices were substracted from the fluorescence value of the stained slices. Results are shown as mean ± SEM.

## 4. Discussion

This study demonstrates an abnormal extracellular nucleotide metabolism in calcific aortic valve disease, which comprises a number of changes in ecto-enzyme activities on a variety of cell types **(Figure 7)**. Consequently, stenotic aortic valves were characterized by reduced levels of extracellular ATP removal and impaired production of adenosine. Moreover, already reduced levels of extracellular adenosine were immediately degraded further due to elevated rate of adenosine deamination.

**Figure 7.**
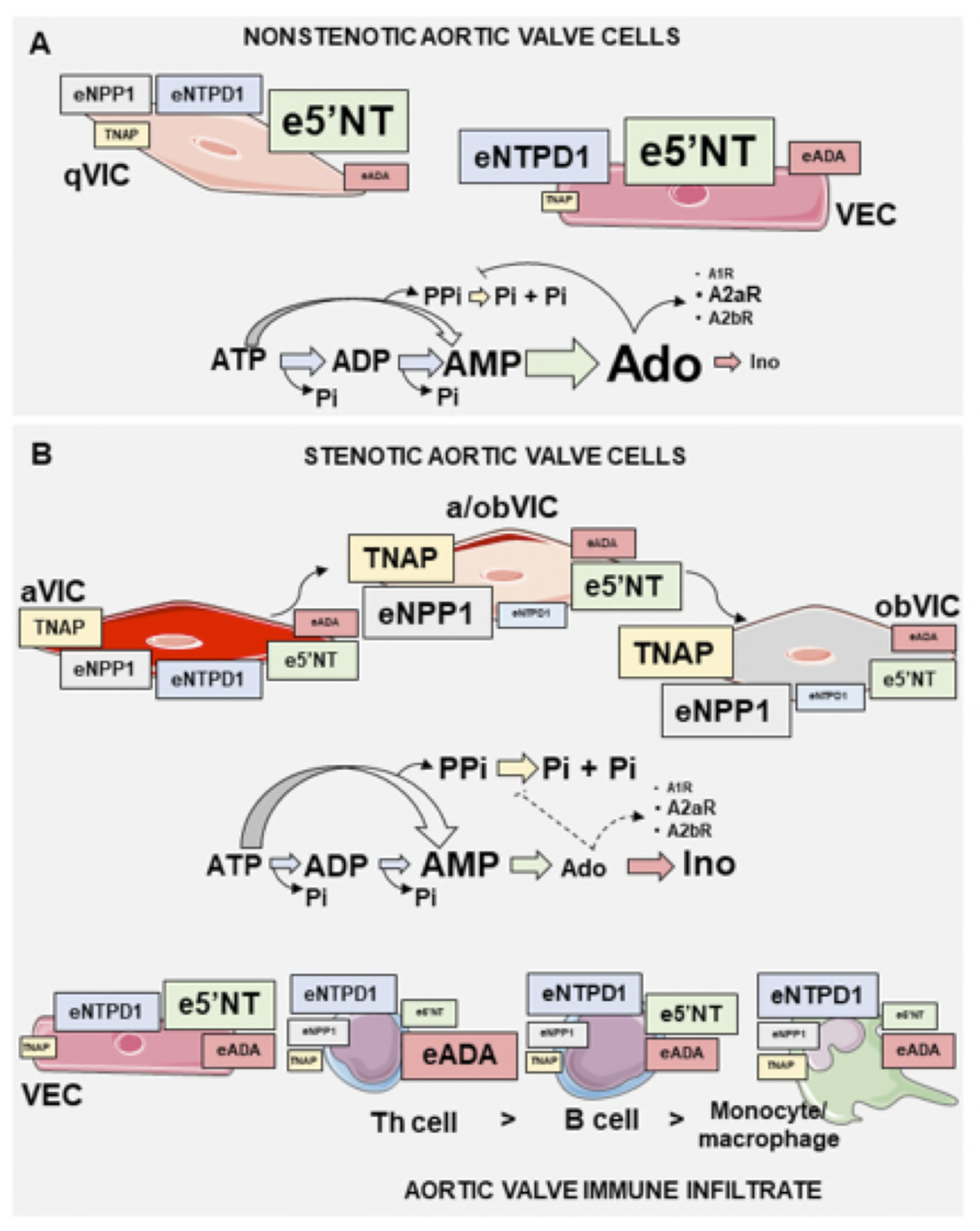
Schematic pathway of extracellular nucleotide and adenosine metabolism in calcific aortic valve disease. In nonstenotic aortic valve **(A)**, ecto-nucleotidases including ecto-nucleoside triphosphate diphosphohydrolase (eNTPD1), ecto-nucleotide pyrophosphatase/ phosphodiesterase 1 (eNPP1) and ecto-5’nucleotidase efficiently produce adenosine on the surface of valvular endothelial cells (VEC) and quiescent valvular interstitial cells (qVIC). These cells exhibit low activities of tissue non-specific alkaline phosphatase (TNAP), which is inhibited by adenosine. In stenotic aortic valve **(B)**, valvular endothelial cells undergo transformation from activated VIC (aVIC) via the transient phenotype (a/obVIC) to osteoblast-like VIC (obVIC). It is related to increased activity of eNPP1 and TNAP, as well as decreased activities of eNTPD1 and e5’NT, that may result in a decrease in the bioavailability of extracellular adenosine and increased degradation of pyrophosphate (PPi) to inorganic orthophosphate (Pi). Stenotic aortic valve immune infiltrate, which mainly consists of T helper cells (Th), B cells and monocytes/macrophages, is a minor source of nucleotide degrading ecto-nucleotidases and a major source of ecto-adenosine deaminase. Therefore in stenotic aortic valve, there is an ecto-enzyme pattern that affect nucleotide and particularly adenosine concentrations to favor a pro-inflammatory milieu, augmenting valve calcification.

For the first time, we thoroughly analyzed the entire aortic valve surfaces and revealed that above metabolic pattern was observed only on the fibrosa surface of stenotic aortic valve and could favor a pro-inflammatory and pro-thrombotic nucleotide milieu and reduction of protective adenosine. (43,44) This is consistent with the pathology and mineral deposition on the aortic side of the valve (fibrosa), where turbulent blood flow contributes to the endothelial disruption and blood retention during valve closure. (45)

Substrates for extracellular enzymes may be released by different cells in the entire circulation, including stimulated cells localized within the aortic valve. (9) Also, availability of particular nucleotide catabolism ecto-enzymes is variable and depends on cell type and each cell’s specific functions. In the cardiovascular system, the most important role in the extracellular ATP catabolism is attributed to the family of ecto-nucleoside triphosphate diphosphohydrolases (eNTPDases). As it has been shown so far, the major member of this family, eNTPD1/CD39 is predominantly expressed in the vasculature by endothelial cells and vascular smooth muscle cells (VSMC). (46) Another enzyme involved in extracellular ATP degradation is ecto-nucleotide pyrophosphatase/ phosphodiesterase 1 (eNPP1) (18) that has been found at high levels in valvular interstitial cells during CAVD. (19) In our study, we confirmed the presence of eNPP1 in aortic valves by immunofluorescence and found its activity on the surface of hVIC and to some extent on inflammatory cells that can infiltrate stenotic aortic valve. However, these cells and, above all, valvular endothelial cells expressed also eNTPD1 in our immunofluorescence, flow cytometry and biochemical studies. Considering previously described increase in eNPP1 expression in CAVD (19), we assume that the decreased total ATP hydrolysis on the fibrosa surface of stenotic valve is the effect of diminished eNTPD1 activity, which expression was reduced in stenotic valves. In recent studies, we have shown the decreased activity and protein level of eNTPD1 in the homogenates of stenotic aortic valves using functional assays, immunohistochemistry (47) and proteomics (48). In addition, in this work we demonstrated a lower level of eNTPD1 on hVIC ongoing differentiation into osteoblast-like cells whose marker was bone sialoprotein. Based on controlling extracellular purinergic gradient, the reduction in eNTPD1 activity can have a number of consequences in CAVD development, particularly associated with inflammation and thrombosis. It has been shown that systemic administration of eNTPD1 minimized injury-induced platelet deposition and leukocyte recruitment, (49,50) while CD39 knockout mice decreased neointimal formation associated with impaired VSMC migration. (16) Moreover, the decreased eNTPD1 activity promotes extracellular ATP accumulation that could stimulate hVIC calcification via P2 purinergic receptor activation. (7)

Although these effects were evoked by the depletion of the extracellular ATP pool, they could be expressed even more strongly through the cooperation of eNTPD1 with adenosine-producing e5NT activity. e5NT was found in a variety of tissues, including abundant activity in vascular endothelium. (51) We have demonstrated that e5NT was the most important ecto-enzyme responsible for AMP hydrolysis on the surface of valvular endothelial and interstitial cells, as well as on the immune cells isolated from stenotic aortic valves. However, the total rate of AMP to adenosine hydrolysis and hence e5NT activity was about 100 times higher on valve cells than on immune cells isolated from stenotic valves. Our immunohistochemical (47) and current immunofluorescence data revealed that a part of the signal for e5NT can accumulate in the areas of calcification, while in not calcified sections of stenotic aortic valves, we observed rather weak signal for this protein and its reduced activity in comparison to non-stenotic valves. We also demonstrated decreased activity (47) and expression (current study) of e5NT throughout the entire valve. In contrast to our results, it has been reported previously that stenotic aortic valves revealed overexpression of e5NT. (8) However, the analysis of only selected fragment of the stenotic valve could be overestimated due to the accumulation of e5NT protein within calcifications.

e5NT-derived adenosine performs many critical functions in the vasculature including suppression of tissue-nonspecific alkaline phosphatase (TNAP), an important enzyme in regulating intracellular calcification. (52) Inorganic pyrophosphate (PPi) that is a substrate for TNAP, is produced by valvular interstitial cells via eNPP1 activity and it is considered as a potent inhibitor of calcification. (53) *Ex vivo* models of aortic valve calcification showed that pig valvular leaflets were stimulated to calcify by the degradation of PPi to the inorganic phosphate (Pi), an inductor of calcification, through the activity of TNAP. (54) We also demonstrated that immunofluorescence signal for both proteins, alkaline phosphatase and eNPP1 was abundant in stenotic aortic valves, with significant accumulation of alkaline phosphatase within the calcification areas.

Patients with mutations in e5NT gene exhibited ectopic calcification within the cardiovascular system. (55) These reports suggest that inhibitors of TNAP, like adenosine, could be considered as potential preventive strategies in CAVD. (56)

However, these beneficial adenosine effects could be also dependent on the type of activated adenosine receptors and the activity of endothelial or immune cell-surface ecto-adenosine deaminase. It has been shown that the stimulation of A1 receptors (A1R) promoted an anti-mineralizing response, whereas A2a receptor (A2aR) activation had the opposite effect. (8) In our study, A2aR was the most abundant in aortic valves but its expression was lower in stenotic valves than in nonstenotic that may be a mechanism for compensating a further development of CAVD. Moreover, we demonstrated that except A1R and A2aR also A2b adenosine receptor (A2bR) was present in human aortic valves. Stimulation of A2bR can cause many positive outcomes, including endothelial protection (57), lipid-lowering (58) and anti-inflammatory (59) effects. It is also essential that A2bR is activated only by high adenosine concentration (micromolar) and therefore A2bR-dependent effects will be triggered when the production of adenosine is maintained by ecto-nucleotidases and it is not excessively degraded by eADA. (60)

Lymphocytes and monocytes/macrophages isolated from stenotic aortic valves were characterized by high expression of ADA-binding protein. Whereas, functional assays using these cells revealed high activity of eADA, which is a key regulator of their function. (61,62) Therefore, increased activity of eADA could be related to the severity of immune infiltration in stenotic aortic valves, which was mainly consisted from CD4+ T cells, CD19+ B cells and CD11b^+^CD14^+^ monocytes/macrophages. However, eADA activity could also be a reflection of endothelial activation, since such pathological conditions as hypoxia, inflammation and atherogenic lipoproteins enhance its endothelial activity what drives vascular and valvular damage. (26,63,64)

In the face of confusing reports about ecto-nucleotidases and adenosine signaling in CAVD, it is crucial to emphasise that complex purinergic signaling pathways involve the deregulation of many ecto-enzyme activities and adenosine receptor expression, which originate from various types of cells that build and pathologically infiltrate aortic valves. Therefore, the wide-spectrum approach should be used both for the analysis of purinergic signaling in CAVD and for the study of potential therapeutic effects of drugs regulating the extracellular pathways of nucleotide and adenosine metabolism. It is clear that enzymes engaged in the extracellular nucleotide cascade might play a significant role in all stages of CAVD, from controlling endothelial damage, through leukocyte infiltration, accumulation of foam cells and secretion of pro-inflammatory mediators to osteoblastic differentiation. Hence, adequate activities of nucleotide and adenosine-regulating ecto-enzymes can be viewed as specific “switches” that shift ATP-driven valvular dysfunction and degeneration toward the states mediated by adenosine, which in turn are dependent on activated adenosine receptors.

## 5. Author contribution

B.K.Z. conceived and conducted the study, performed data analysis and interpretation, and wrote the manuscript. P.J. assisted with material collection, enzymatic assays and data analysis. M.S. and R.B. performed the analysis of mRNA expression. A.B. performed enzymatic assays in PBMC. P.Z. assisted with determination of valve deposits compounds concentrations. D.F. and C.A. assisted in flow cytometry analysis.

A.J. assisted histological and analysis. R.L. and J.R. provided postoperative material.

E.M.S, S.H and J.S. assisted in data analysis and interpretation. M. H. Y. and R.T.S were responsible for concenption and design, final manuscript approval, conceived and conducted the study, and wrote the paper.

## 8.

### Abbreviations

α-SMA: smooth muscle cells alpha actin
A1R: adenosine A1 receptor
A2aR: adenosine A2a receptor
A2bR: adenosine A2b receptor
A3R: adenosine A3 receptor
ADP: adenosine diphosphate
AMP: adenosine monophosphate
AOPCP: 5’-(α,β-methylene)diphosphate
AP: alkaline phosphatase
ARL67156: 6-N,N-Diethyl-β-γ-dibromomethylene-D-adenosine-5’-triphosphate trisodium salt hydrate
ATP: adenosine triphosphate
aVIC: activated valvular interstitial cells
AVR: aortic valve replacement
CAVD: calcific aortic valve disease
e5NT: ecto-5’nucleotidase
eADA: ecto-adenosine deaminase
EHNA: erythro-9-(2-hydroxy-3-nonyl)adenine
eNPP1: ecto-nucleotide pyrophosphatase/phosphodiesterase 1
eNPPs: ecto-nucleotide pyrophosphatases/ phosphodiesterases
eNTPD1: ecto-nucleoside triphosphate diphosphohydrolase 1
FBS: fetal bovine serum
HBSS: Hanks Balanced Salt Solution
HE: hematoxylin and eosin staining
HPLC: high performance liquid chromatography
VEC: aortic valve endothelial cells
hVIC: aortic valve interstitial cells
LDL: low density lipoproteins
NBTI: S-(4-Nitrobenzyl)-6-thioinosine
obVIC: osteoblast-like valvular interstitial cells
OMSB: Orcein Martius Scarlet Blue staining
PBMC: peripheral blood mononuclear cells
PBS: phosphate buffered saline
Pi: inorganic phosphate
PPADS: pyridoxal phosphate-6-azo(benzene-2,4-disulfonic acid) tetrasodium salt hydrate
PPi: inorganic pyrophosphate
qVIC: quiescent valvular interstitial cells
TAVI: transcatheter aortic valve implantation
TNAP: tissue nonspecific alkaline phosphatase
TR: Masson’s Trichrome staining
VSMC: vascular smooth muscle cells
vWF: von Wilebrant factor

## References

1. Dweck MR, Boon NA, Newby DE. Calcific aortic stenosis: a disease of the valve and the myocardium. J Am Coll Cardiol. 2012;60(19):1854–63.

2. Bonow RO, Carabello BA, Chatterjee K, de Leon AC, Faxon DP, Freed MD, et al. ACC/AHA 2006 practice guidelines for the management of patients with valvular heart disease: executive summary. J Am Coll Cardiol. 2006;48(3):598–675.

3. Beckmann E, Grau JB, Sainger R, Poggio P, Ferrari G. Insights into the use of biomarkers in calcific aortic valve disease. J Heart Valve Dis. 2010;19(4):441.

4. Mohler Iii ER. Mechanisms of aortic valve calcification. Am J Cardiol. 2004;94(11):1396–402.

5. Liu AC, Joag VR, Gotlieb AI. The emerging role of valve interstitial cell phenotypes in regulating heart valve pathobiology. Am J Pathol. 2007;171(5):1407–18.

6. Freeman R V, Otto CM. Spectrum of calcific aortic valve disease pathogenesis, disease progression, and treatment strategies. Circulation. 2005;111(24):3316–26.

7. Osman L, Chester AH, Amrani M, Yacoub MH, Smolenski RT. A novel role of extracellular nucleotides in valve calcification - A potential target for atorvastatin. Circulation [Internet]. 2006;114:I566–72. Available from: %3CGo

8. Mahmut A, Boulanger MC, Bouchareb R, Hadji F, Mathieu P. Adenosine derived from ecto-nucleotidases in calcific aortic valve disease promotes mineralization through A2a adenosine receptor. Cardiovasc Res. 2015;

9. Erlinge D, Burnstock G. P2 receptors in cardiovascular regulation and disease. Purinergic Signal. 2008;4(1):1–20.

10. Yegutkin GG, Mikhailov A, Samburski SS, Jalkanen S. The detection of micromolar pericellular ATP pool on lymphocyte surface by using lymphoid ecto-adenylate kinase as intrinsic ATP sensor. Mol Biol Cell. 2. 2006;17(8):3378–85.

11. Sumi Y, Woehrle T, Chen Y, Bao Y, Li X, Yao Y, et al. Plasma ATP is required for neutrophil activation in a mouse sepsis model. Shock. 2014;

12. Bours MJL, Swennen ELR, Di Virgilio F, Cronstein BN, Dagnelie PC. Adenosine 5 ‘-triphosphate and adenosine as endogenous signaling molecules in immunity and inflammation. Pharmacol Ther [Internet]. 2006;112(2):358–404. Available from: %3CGo

13. Atkinson B, Dwyer K, Enjyoji K, Robson SC. Ecto-nucleotidases of the CD39/NTPDase family modulate platelet activation and thrombus formation: Potential as therapeutic targets. Blood Cells Mol Dis. 2006;36(2):217–22.

14. Hoebertz A, Arnett TR, Burnstock G. Regulation of bone resorption and formation by purines and pyrimidines. Trends Pharmacol Sci. 2003;24(6):290–7.

15. Kaniewska E, Sielicka A, Sarathchandra P, Pelikant-Malecka I, Olkowicz M, Slominska EM, et al. Immunohistochemical and functional analysis of ectonucleoside triphosphate diphosphohydrolase 1 (CD39) and ecto-5’-nucleotidase (CD73) in pig aortic valves. Nucleosides Nucleotides Nucleic Acids. 2014;33(4–6):305–12.

16. Behdad A, Sun X, Khalpey Z, Enjyoji K, Wink M, Wu Y, et al. Vascular smooth muscle cell expression of ectonucleotidase CD39 (ENTPD1) is required for neointimal formation in mice. Purinergic Signal. 2009;5(3):335–42.

17. Deaglio S, Dwyer KM, Gao W, Friedman D, Usheva A, Erat A, et al. Adenosine generation catalyzed by CD39 and CD73 expressed on regulatory T cells mediates immune suppression. J Exp Med. 2007;204(6):1257–65.

18. Yegutkin GG. Nucleotide- and nucleoside-converting ectoenzymes: Important modulators of purinergic signalling cascade. Biochim Biophys Acta-Molecular Cell Res [Internet]. 2008;1783(5):673–94. Available from: %3CGo

19. Côté N, El Husseini D, Pépin A, Guauque-Olarte S, Ducharme V, Bouchard-Cannon P, et al. ATP acts as a survival signal and prevents the mineralization of aortic valve. J Mol Cell Cardiol. 2012;52(5):1191–202.

20. Mathieu P, Boulanger MC, Bouchareb R. Molecular biology of calcific aortic valve disease: towards new pharmacological therapies. Expert Rev Cardiovasc Ther. 2014;12(7):851–62.

21. Toczek M, Kutryb-Zajac B, Kapczynska M, Lipinski M, Slominska EM, Smolenski RT. Extracellular adenine nucleotide catabolism in heart valves. Nucleosides Nucleotides Nucleic Acids. 2014;33(4–6):329–32.

22. Reiss AB, Cronstein BN. Regulation of foam cells by adenosine. Arter Thromb Vasc Biol. 2012;32(4):879–86.

23. Eltzschig HK. Adenosine: an old drug newly discovered. Anesthesiology. 2009;111(4):904–15.

24. Antonioli L, Csoka B, Fornai M, Colucci R, Kokai E, Blandizzi C, et al. Adenosine and inflammation: what’s new on the horizon? Drug Discov Today. 2014;19(8):1051–68.

25. Orriss IR, Burnstock G, Arnett TR. Purinergic signalling and bone remodelling. Curr Opin Pharmacol. 2010;10(3):322–30.

26. Kutryb-Zajac B, Mateuszuk L, Zukowska P, Jasztal A, Zabielska M, Toczek M, et al. Increased activity of vascular adenosine deaminase in atherosclerosis and therapeutic potential of its inhibition. Cardiovasc Res. 2016;112:590–605.

27. Henderson JF, Brox L, Zombor G, Hunting D, Lomax CA. Specificity of adenosine deaminase inhibitors. Biochem Pharmacol [Internet]. 1977 Nov 1 [cited 2018 Aug 11];26(21):1967–72. Available from: https://www.sciencedirect.com/science/article/abs/pii/000629527790003X

28. Plagemann PG, Wohlhueter RM. Effects of nucleoside transport inhibitors on the salvage and toxicity of adenosine and deoxyadenosine in L1210 and P388 mouse leukemia cells. Cancer Res [Internet]. 1985 Dec [cited 2018 Aug 11];45(12 Pt 1):6418–24. Available from: http://www.ncbi.nlm.nih.gov/pubmed/3877568

29. Smolenski RT, Lachno DR, Ledingham SJM, Yacoub MH. Determination of sixteen nucleotides, nucleosides and bases using high-performance liquid chromatography and its application to the study of purine metabolism in hearts for transplantation. J Chromatogr B Biomed Sci Appl [Internet]. 1990 Jan [cited 2016 Sep 12];527:414–20. Available from: http://linkinghub.elsevier.com/retrieve/pii/S0378434700821258

30. Michaylova V, Ilkova P. Photometric determination of micro amounts of calcium with arsenazo III. Anal Chim Acta. 1971;53(1):194–8.

31. Chauhan UPS, Ray Sarkar BC. Use of calmagite for the determination of traces of magnesium in biological materials. Anal Biochem. 1969;32(1):70–80.

32. Feng J, Chen Y, Pu J, Yang X, Zhang C, Zhu S, et al. An improved malachite green assay of phosphate: Mechanism and application. Anal Biochem. 2011;409(1):144–9.

33. Gajda M, Jasztal A, Banasik T, Jasek-Gajda E, Chlopicki S. Combined orcein and martius scarlet blue (OMSB) staining for qualitative and quantitative analyses of atherosclerotic plaques in brachiocephalic arteries in apoE/LDLR−/− mice. Histochem Cell Biol [Internet]. 2017 Jun 6 [cited 2018 Feb 5];147(6):671–81. Available from: http://link.springer.com/10.1007/s00418-017-1538-8

34. Bartoszewski R, Rab A, Fu L, Bartoszewska S, Collawn J, Bebok Z. CFTR Expression Regulation by the Unfolded Protein Response. In: Methods in enzymology [Internet]. 2011 [cited 2018 Feb 6]. p. 3–24. Available from: http://www.ncbi.nlm.nih.gov/pubmed/21329791

35. Bartoszewski R, Hering A, Marszałł M, Stefanowicz Hajduk J, Bartoszewska S, Kapoor N, et al. Mangiferin Has an Additive Effect on the Apoptotic Properties of Hesperidin in Cyclopia sp. Tea Extracts. Trajkovic V, editor. PLoS One [Internet]. 2014 Mar 14 [cited 2018 Feb 6];9(3):e92128. Available from: http://dx.plos.org/10.1371/journal.pone.0092128

36. Larionov A, Krause A, Miller W. A standard curve based method for relative real time PCR data processing. BMC Bioinformatics [Internet]. 2005 Mar 21 [cited 2018 Feb 6];6(1):62. Available from: http://www.ncbi.nlm.nih.gov/pubmed/15780134

37. Bhattarai S, Freundlieb M, Pippel J, Meyer A, Abdelrahman A, Fiene A, et al. α,β-Methylene-ADP (AOPCP) Derivatives and Analogues: Development of Potent and Selective ecto −5′-Nucleotidase (CD73) Inhibitors. J Med Chem [Internet]. 2015 Aug 13 [cited 2018 Aug 11];58(15):6248–63. Available from: http://www.ncbi.nlm.nih.gov/pubmed/26147331

38. Van Belle H. Alkaline phosphatase. I. Kinetics and inhibition by levamisole of purified isoenzymes from humans. Clin Chem [Internet]. 1976 Jul [cited 2018 Aug 11];22(7):972–6. Available from: http://www.ncbi.nlm.nih.gov/pubmed/6169

39. Crack BE, Pollard CE, Beukers MW, Roberts SM, Hunt SF, Ingall AH, et al. Pharmacological and biochemical analysis of FPL 67156, a novel, selective inhibitor of ecto-ATPase. Br J Pharmacol [Internet]. 1995 Jan [cited 2018 Aug 11];114(2):475–81. Available from: http://www.ncbi.nlm.nih.gov/pubmed/7533620

40. Yegutkin GG, Wieringa B, Robson SC, Jalkanen S. Metabolism of circulating ADP in the bloodstream is mediated via integrated actions of soluble adenylate kinase-1 and NTPDase1/CD39 activities. Faseb j. 2012;26(9):3875–83.

41. Vollmayer P, Clair T, Goding JW, Sano K, Servos J, Zimmermann H. Hydrolysis of diadenosine polyphosphates by nucleotide pyrophosphatases/phosphodiesterases. Eur J Biochem [Internet]. 2003 Jul [cited 2018 Aug 11];270(14):2971–8. Available from: http://www.ncbi.nlm.nih.gov/pubmed/12846830

42. Kleiveland CR. Peripheral Blood Mononuclear Cells. In: The Impact of Food Bioactives on Health [Internet]. Cham: Springer International Publishing; 2015 [cited 2018 Feb 6]. p. 161–7. Available from: http://link.springer.com/10.1007/978-3-319-16104-4_15

43. Haskó G, Cronstein B. Regulation of Inflammation by Adenosine. Front Immunol [Internet]. 2013 [cited 2018 Aug 13];4:85. Available from: http://www.ncbi.nlm.nih.gov/pubmed/23580000

44. Johnston-Cox H, Ravid K. Adenosine and blood platelets. Purinergic Signal [Internet]. 2011;7(3):357–65. Available from: http://dx.doi.org/10.1007/s11302-011-9220-4

45. Yutzey KE, Demer LL, Body SC, Huggins GS, Towler DA, Giachelli CM, et al. Calcific Aortic Valve Disease A Consensus Summary From the Alliance of Investigators on Calcific Aortic Valve Disease. Arterioscler Thromb Vasc Biol. 2014;ATVBAHA-114.

46. Kauffenstein G, Drouin A, Thorin-Trescases N, Bachelard H, Robaye B, D’Orleans-Juste P, et al. NTPDase1 (CD39) controls nucleotide-dependent vasoconstriction in mouse. Cardiovasc Res. 2010;85(1):204–13.

47. Kaniewska-Bednarczuk E, Kutryb-Zajac B, Sarathchandra P, Pelikant-Malecka I, Sielicka A, Piotrowska I, et al. CD39 and CD73 in the aortic valve— biochemical and immunohistochemical analysis in valve cell populations and its changes in valve mineralization. Cardiovasc Pathol [Internet]. 2018 Sep 7 [cited 2018 Aug 13];36:53–63. Available from: http://www.ncbi.nlm.nih.gov/pubmed/30056298

48. Olkowicz M, Jablonska P, Rogowski J, Smolenski RT. Simultaneous accurate quantification of HO-1, CD39, and CD73 in human calcified aortic valves using multiple enzyme digestion – filter aided sample pretreatment (MED-FASP) method and targeted proteomics. Talanta [Internet]. 2018 May 15 [cited 2018 Aug 13];182:492–9. Available from: http://www.ncbi.nlm.nih.gov/pubmed/29501184

49. Drosopoulos JH, Kraemer R, Shen H, Upmacis RK, Marcus AJ, Musi E. Human solCD39 inhibits injury-induced development of neointimal hyperplasia. Thromb Haemost. 2010;103(2):426–34.

50. Robson SC, Wu Y, Sun X, Knosalla C, Dwyer K, Enjyoji K. Ectonucleotidases of CD39 family modulate vascular inflammation and thrombosis in transplantation. Semin Thromb Hemost. 2005;31(2):217–33.

51. Thompson LF, Eltzschig HK, Ibla JC, Van De Wiele CJ, Resta R, Morote-Garcia JC, et al. Crucial role for ecto-5’-nucleotidase (CD73) in vascular leakage during hypoxia. J Exp Med. 2004;200(11):1395–405.

52. Gan XT, Taniai S, Zhao G, Huang CX, Velenosi TJ, Xue J, et al. CD73-TNAP crosstalk regulates the hypertrophic response and cardiomyocyte calcification due to alpha1 adrenoceptor activation. Mol Cell Biochem. 2014;394(1–2):237–46.

53. Towler DA. Inorganic pyrophosphate: a paracrine regulator of vascular calcification and smooth muscle phenotype. Arter Thromb Vasc Biol. 2005;25:651–4.

54. Swetha R, Ajit Y, Charles O. The Role of Inorganic Pyrophosphate in Aortic Valve Calcification. J Heart Valve Dis. 2014;23:387–94.

55. St Hilaire C, Ziegler SG, Markello TC, Brusco A, Groden C, Gill F, et al. NT5E mutations and arterial calcifications. N Engl J Med. 2011;364(5):432–42.

56. Jin H, St Hilaire C, Huang Y, Yang D, Dmitrieva NI, Negro A, et al. Increased activity of TNAP compensates for reduced adenosine production and promotes ectopic calcification in the genetic disease ACDC. Sci Signal [Internet]. 2016 Dec 13 [cited 2018 Feb 5];9(458):ra121. Available from: http://www.ncbi.nlm.nih.gov/pubmed/27965423

57. Eckle T, Faigle M, Grenz A, Laucher S, Thompson LF, Eltzschig HK. A2B adenosine receptor dampens hypoxia-induced vascular leak Running Title: A2BAR in vascular permeability. 2007 [cited 2018 Feb 5]; Available from: http://www.bloodjournal.org/content/bloodjournal/early/2007/12/04/blood-2007-10-117044.full.pdf?sso-checked=true

58. Koupenova M, Johnston-Cox H, Vezeridis A, Gavras H, Yang D, Zannis V, et al. A2b adenosine receptor regulates hyperlipidemia and atherosclerosis. Circulation [Internet]. 2012 Jan 17 [cited 2016 Dec 11];125(2):354–63. Available from: http://www.ncbi.nlm.nih.gov/pubmed/22144568

59. Aherne CM, Kewley EM, Eltzschig HK. The resurgence of A2B adenosine receptor signaling. Biochim Biophys Acta - Biomembr [Internet]. 2011 May 1 [cited 2018 Feb 5];1808(5):1329–39. Available from: https://www.sciencedirect.com/science/article/pii/S0005273610001653#f0015

60. Haskó G, Csóka B, Németh ZH, Vizi ES, Pacher P. A(2B) adenosine receptors in immunity and inflammation. Trends Immunol [Internet]. 2009 Jun [cited 2018 Aug 13];30(6):263–70. Available from: http://www.ncbi.nlm.nih.gov/pubmed/19427267

61. Antonioli L, Colucci R, La Motta C, Tuccori M, Awwad O, Da Settimo F, et al. Adenosine deaminase in the modulation of immune system and its potential as a novel target for treatment of inflammatory disorders. In: Curr Drug Targets. Netherlands; 2012. p. 842–62.

62. Lluis C, Franco R, Cordero O. Ecto-ADA in the development of the immune system. In: Immunol Today. England; 1998. p. 533–4.

63. Eltzschig HK, Faigle M, Knapp S, Karhausen J, Ibla J, Rosenberger P, et al. Endothelial catabolism of extracellular adenosine during hypoxia: the role of surface adenosine deaminase and CD26. Blood. 2006;108(5):1602–10.

64. Kutryb-Zajac B, Zukowska P, Toczek M, Zabielska M, Lipinski M, Rybakowska I, et al. Extracellular Nucleotide Catabolism in Aortoiliac Bifurcation of Atherosclerotic ApoE/LDLr Double Knock Out Mice. Nucleosides Nucleotides Nucleic Acids. 2014;33(4–6):323–8.

